# *In vivo* characterisation of fluorescent proteins in budding yeast

**DOI:** 10.1101/431874

**Authors:** Dennis Botman, Daan Hugo de Groot, Phillipp Schmidt, Joachim Goedhart, Bas Teusink

**Affiliations:** Systems Bioinformatics/AIMMS, Vrije Universiteit Amsterdam, De Boelelaan 1085, 1081 HV Amsterdam, The Netherlands.; Section of Molecular Cytology, van Leeuwenhoek Centre for Advanced Microscopy, Swammerdam Institute for Life Sciences, University of Amsterdam, Science Park 904, 1098 XH Amsterdam, The Netherlands.

## Abstract

Fluorescent proteins (FPs) are widely used in many organisms, but are commonly characterised *in vitro.* However, the *in vitro* properties may poorly reflect *in vivo* performance. Therefore, we characterised 27 FPs *in vivo* using *Saccharomyces cerevisiae* as model organism. We linked the FPs via a T2A peptide to a control FP, producing equimolar expression of the 2 FPs from 1 plasmid. Using this strategy, we characterised the FPs for brightness, photostability, photochromicity and pH-sensitivity, achieving a comprehensive *in vivo* characterisation. Many FPs showed different *in vivo* properties compared to existing *in vitro* data. Additionally, various FPs were photochromic, which affects readouts due to complex bleaching kinetics. Finally, we codon optimized the best performing FPs for optimal expression in yeast, and found that codon-optimization alters FP characteristics. These FPs improve experimental signal readout, opening new experimental possibilities. Our results may guide future studies in yeast that employ fluorescent proteins.

## Introduction

Fluorescent proteins (FPs) have become a widely used tool for many organisms as they are genetically encoded and do not need a cofactor to fluoresce. This enables visualization and measurements of cellular processes in a spatiotemporal and non-invasive manner. Since the discovery of GFP by Prasher and colleagues (Prasher et al., 1992), new FPs have been developed, each with their own traits (Cranfill et al., 2016; Goedhart et al., 2012; Griesbeck et al., 2001; Heim and Tsien, 1996; Kremers et al., 2007, 2006; Lam et al., 2012; Merzlyak et al., 2007; Nagai et al., 2002; Pédelacq et al., 2006; Shaner et al., 2013, 2008, 2004, Yang et al., 1998, 1996). Not a single FP is optimal for all possible experiments, since every FP has its strong and weak characteristics. Based on the specific characteristics needed for an experiment, one should choose the most suitable FP. Various characteristics can be considered.

The two most important characteristics for live cell imaging are brightness and photostability as these determine the strength of the fluorescent signal and its ability to maintain it over time. The *in vitro* brightness is often defined as the multiplication of the quantum yield (the amount of photons emitted per absorbed photons) and the extinction coefficient (the amount of absorbed photons at a specific wavelength). In contrast, the *in vivo* (or practical) brightness also depends on the level of functional, fluorescent FPs which is determined by protein folding, maturation and degradation. Moreover, other factors such as the cellular environment and post-translation modification can affect the practical brightness. Therefore, the *in vitro* brightness is often not directly proportional to the (practical) *in vivo* brightness. For experiments in living organisms, the *in vivo* brightness is more relevant (Bindels et al., 2017; Heppert et al., 2016; Lee et al., 2013).

The loss of fluorescence intensity due to illumination of a fluorophore is known as photobleaching. Upon excitation, electrons can transition from the excited singlet state to the excited triplet state and subsequently interact with other molecules, and this can irreversibly modify and damage the chromophore (Donnert et al., 2007; Lichtman and Conchello, 2005). The number of excitation and emission cycles an FP can undergo before it bleaches depends on the specific FP and the illumination settings (Cranfill et al., 2016). A photostable FP is obviously desired for live cell imaging. Next to that, simple bleaching kinetics (i.e. mono exponential decay) are desirable as this simplifies corrections or predictions for bleaching.

Yet, bleaching kinetics can be complex due to reversible bleaching processes, for which we use the term photochromism. This under-appreciated process is related to bleaching and is caused by a reversible dark state of the FP chomophore (Bindels et al., 2017; Dean et al., 2011; Dickson et al., 1997; Shaner et al., 2008; Sinnecker et al., 2005). Photochromism results in reversible bleaching when using multiple excitation wavelengths that give complex and unpredictable bleaching kinetics. This bleaching is called reversible because the FPs can transit from the reversible dark state back to the fluorescent state, which increases the fluorescent signal in time. The mechanisms underlying photochromism are probably due to a cis-trans conversion of the chromophore tyrosyl side chain together with a protonation of the chromophore (Andresen et al., 2005; Henderson et al., 2007; Mizuno et al., 2008; Pletnev et al., 2008). Although photochromism can greatly affect readouts in multicolour timelapse experiments, it has hardly been systematically characterised, and the effect of each different excitation wavelength on photochromism is poorly documented. Therefore, the photochromism of FPs is largely unknown and characterisation is needed, ideally *in vivo.*

Fourth, monomeric behaviour of FPs is important as they have a natural tendency to form dimers or oligomers, which affects localisation and the functionality of tagged proteins (Baird et al., 2000; Tsien, 1998). Optimization of *Aequorea victoria* derived FPs has led to monomeric variants in which the hydrophobic amino acids at the dimer interface (i.e. Ala^206^, Leu^221^ and Phe^223^) have been replaced with positive charged amino acids (i.e. A206K, L221K, or F223R) (Zacharias et al., 2002). These mutations are specific for FPs derived from *Aequorea victoria,* and similar mutations in other FPs may not work. The generation of monomeric FPs from obligate tetramers requires substantial engineering, which usually is accompanied by a loss in brightness (Baird et al., 2000; Campbell et al., 2002; Shaner et al., 2004). Oligomerisation tendency has been extensively characterised *in vivo* in mammalian cells but it is not known whether these results hold in other species (Cranfill et al., 2016).

Fifth, pH sensitivity determines how much a pH change affects the fluorescence of FPs. FPs with low pH sensitivity are necessary for experiments in acidic environments or under conditions of dynamic pH changes, something that is very common in yeast (Orij et al., 2012). Quenching by pH occurs through the protonation and deprotonation of the chromophore side chains of FPs (Shinoda et al., 2018). The pH quenching curves can best be described by a Hill fit that gives a pKa value and a Hill coefficient. The pH robustness is often misinterpreted by only looking at pKa values. For instance, an FP with a low pKa and a low Hill coefficient does not show a pH range in which the fluorescence remains constant. Therefore, these FPs are still pH-sensitive, even with a low pKa. On the other hand, an FP with a low pKa and a high Hill coefficient is pH-insensitive as this FP has a plateau at pH values above the pKa. Thus, pH-insensitive FPs are better identified based on these two parameters combined. In addition, pH sensitivity has always been described *in vitro,* neglecting the effect of cytolosic components on the pH quenching. How representative *in vitro* pH sensitivity is for the *in vivo* performance is unknown.

Finally, after translation, an FP folds and undergoes various autocatalytic steps before becoming fluorescent, a process called maturation (Balleza et al., 2018; Chudakov et al., 2010; Heim et al., 1994; Reid and Flynn, 1997). Although FPs do not need a cofactor for maturation, they do need oxygen, an important sidenote for studies in anaerobic conditions. The maturation process determines how fast and how much of newly synthesised FPs will become fluorescent, essential when measuring fluorescence changes in time. Besides, maturation is important for brightness as it determines the percentage of fluorescent proteins and reduces day-to-day variation when it matures reliably.

Clearly, all aforementioned properties of FPs can influence experimental success and reliability of the results. Accordingly, these traits should be characterised systematically to assess which FPs are suitable for an experiment. Until now, FPs are selected for use in yeast or other organisms based on their characteristics in bacterial, mammalian or *in vitro* environments. However, these data may not represent the *in vivo* behaviour of FPs (Bindels et al., 2017; Heppert et al., 2016; Lee et al., 2013). This is explained by the fact that the host organism can change FP traits through post-translational modifications, the cellular environment, protein translation and folding efficiency or growth conditions such as temperature. Since *in vivo* characterisation could help for selecting the most suitable FP for an experiment, we aimed to characterise the mentioned characteristics systemetically *in vivo.*

In the present paper we characterised the mostly commonly used FPs *in vivo* using *Saccharomyces Cerevisiae* as model organism. This is a yeast species that grows optimally at 30°C. By using a variety of assays, we measured FP properties *in vivo.* Accordingly, the presented characterisation reflects the real behaviour of FPs in yeast cells. We found many critical differences between our characterisation *in vivo* in yeast compared to characterisations done in mammalian cell systems or *in vitro.* Furthermore, we codon-optimized the best performing FPs in each spectral class and generated yeast FPs (yFPs). The generated yFPs outperform the conventional FPs and are recommended for future experiments in yeast.

## Material and Methods

### Creation of constructs

sYFP2-T2A-mTq2, tagRFPT-T2A-mTq2, tdTomato-T2A-mTq2, tagRFP-T2A-mTq2, mCherry-T2A-mTq2, mCherry-T2A-eGFP, mCherry-T2A-sGFP2, YPET-T2A-mTq2, mCitrine-T2A-mTq2, mNeongreen-T2A-mTq2, mKOk-T2A-mTq2, mClover-T2A-mTq2, mRuby2-T2A-mTq2, mScarlet-T2A-mTq2, mScarletI-T2A-mTq2, mCherry-T2A-mTq2 and mKate2-T2A-mTq2 were based on mKO2-T2A-mTq2 (addgene plasmid #98838). The plasmids mVenus-mTq2, eCFP and mTFP in a clontech style C1 mammalian expression vector and mCherry in a clontech style N1 mammalian expression vector were made by restriction enzyme based cloning.

pFA6a-link-yoSuperfolderGFP-CaURA3 (Addgene plasmid #44873), pFA6a-link-yomCherry-CaURA3 (Addgene plasmid #44876), pFA6a-link-yoTagRFP-T-CaURA3 (Addgene plasmid #44877), pFA6a-link-yoEGFP-CaURA3 (Addgene plasmid #44872) were a gift from Wendell Lim & Kurt Thorn. pKT90 (pFA6a–link–yEVenus–SpHIS5, Addgene plasmid #8714) and pKT174 (pFA6a–link–yECFP–CaURA3, Addgene plasmid #8720, named yoeCFP in this study) were a gift from Kurt Thorn.

The yeast expression vector pDRF1-GW (a gift from Wolf Frommer & Dominique Loque, Addgene plasmid #36026) with the Nhel and Notl restriction sites was created using the GateWay Kit (Thermo Fisher Scientific, Waltham, MA, USA).

sYFP2-T2A-mTq2, tagRFPT-T2A-mTq2, tdTomato-T2A-mTq2, tagRFP-T2A-mTq2, mCherry-T2A-mTq2, YPET-T2A-mTq2, mCitrine-T2A-mTq2, mNeongreen-T2A-mTq2, mKO2-T2A-mTq2, mKOk-T2A-mTq2, mClover-T2A-mTq2, mRuby2-T2A-mTq2, mScarlet-T2A-mTq2, mScarletI-T2A-mTq2, mCherry-T2A-mTq2 and mKate2-T2A-mTq2 were digested using NheI and NotI (New England Biolabs, Ipswich, Massachusetts, USA) and ligated into pDRF1 diggested with the same enzymes using T4 ligase (New England Biolabs) which created pDRF1 containing sYFP2, mVenus, tagRFP, tagRFPT, mCherry, mCitrine, mNeongreen, YPET, mKO2, mKOk, mClover, mRuby2, tdTomato, mKate2, mCherry, mScarlet and mScarletI fused with T2A-mTq2 in pDRF1.

mVenus-T2A-mTq2 was created by digesting mVenus-mTq2 C1 with NheI and Kpn2I. Next, the digested fragment was ligated using T4 ligase in mCherry-T2A-mTq2 digested with the same enzymes, replacing mCherry with mVenus.

sYFP2-T2A-mCherry C1 was created by digesting mCherry N1 with NotI and BamHI (New England Biolabs). Next, the digested fragment was ligated using T4 ligase in sYFP2-T2A-mTq2 C1 digested with the same enymes, replacing mTq2 for mCherry.

A PCR using KOD polymerase (Merck-Millipore, Burlington, Massachusetts, USA), was performed on yosfGFP-T2A-mCherry, eGFP-T2A-mCherry, yoeGFP-T2A-mCherry and sGFP2-T2A-mCherry in a C1 vector according to table 1. The products and FP-T2A-mTq2 pDRF1 were digested with NheI and Kpn2I (Thermo Fisher Scientific, Waltham, Massachusetts, USA) and the products were ligated into the plasmid, replacing the FP N-terminally of T2A-mTq2 with yosfGFP, eGFP, yoeGFP and sGFP2. Next, mTq2 was cut out of eGFP-T2A-mTq2, mNeongreen-T2A-mTq2, sGFP2-T2A-mTq2, yosfGFP-T2A-mTq2 and yoeGFP-T2A-mTq2 in pDRF1 using Kpn2I and NotI and replaced by mCherry digested from sYFP2-T2A-mCherry using the same enzymes.

**Table 1.**
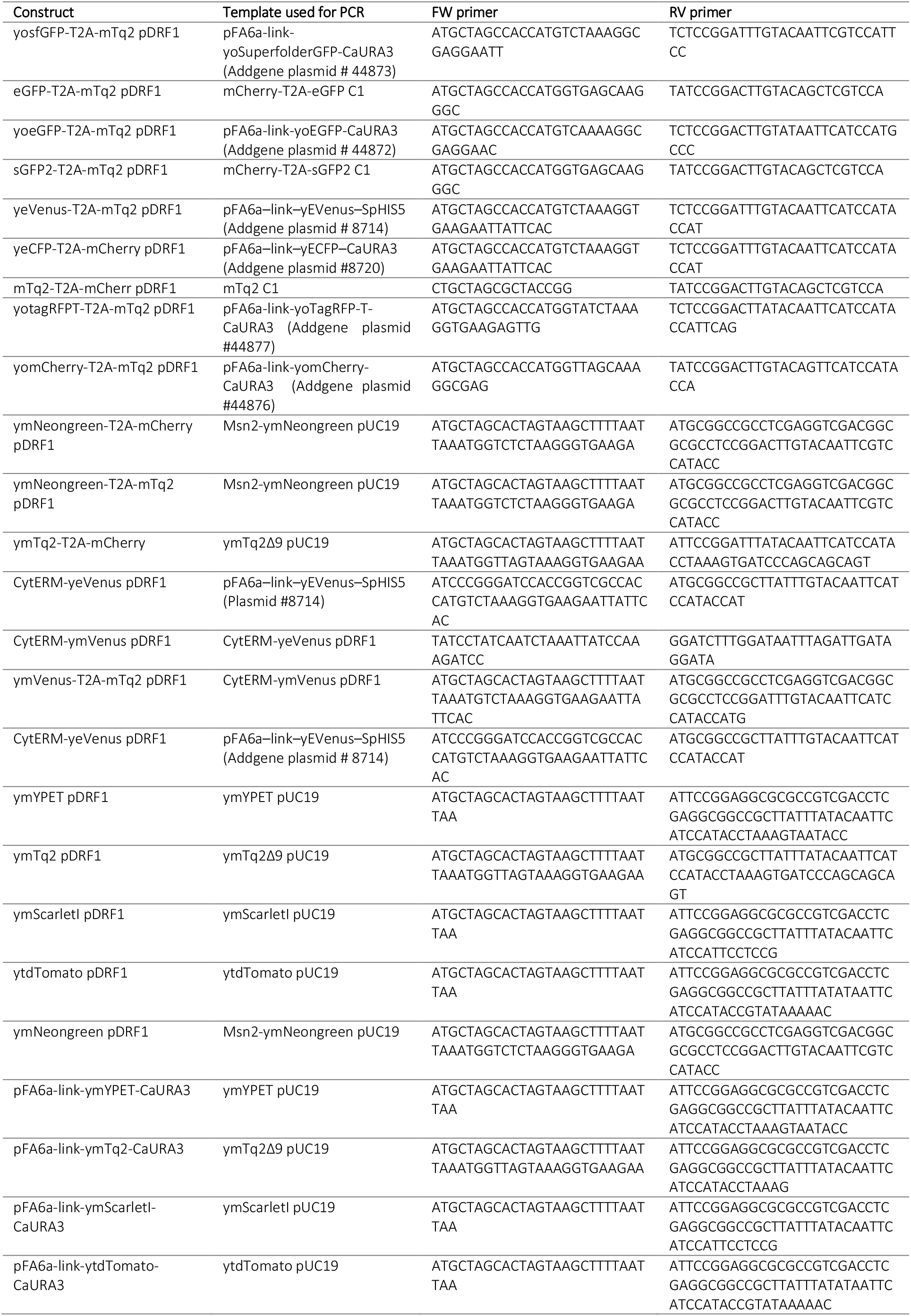
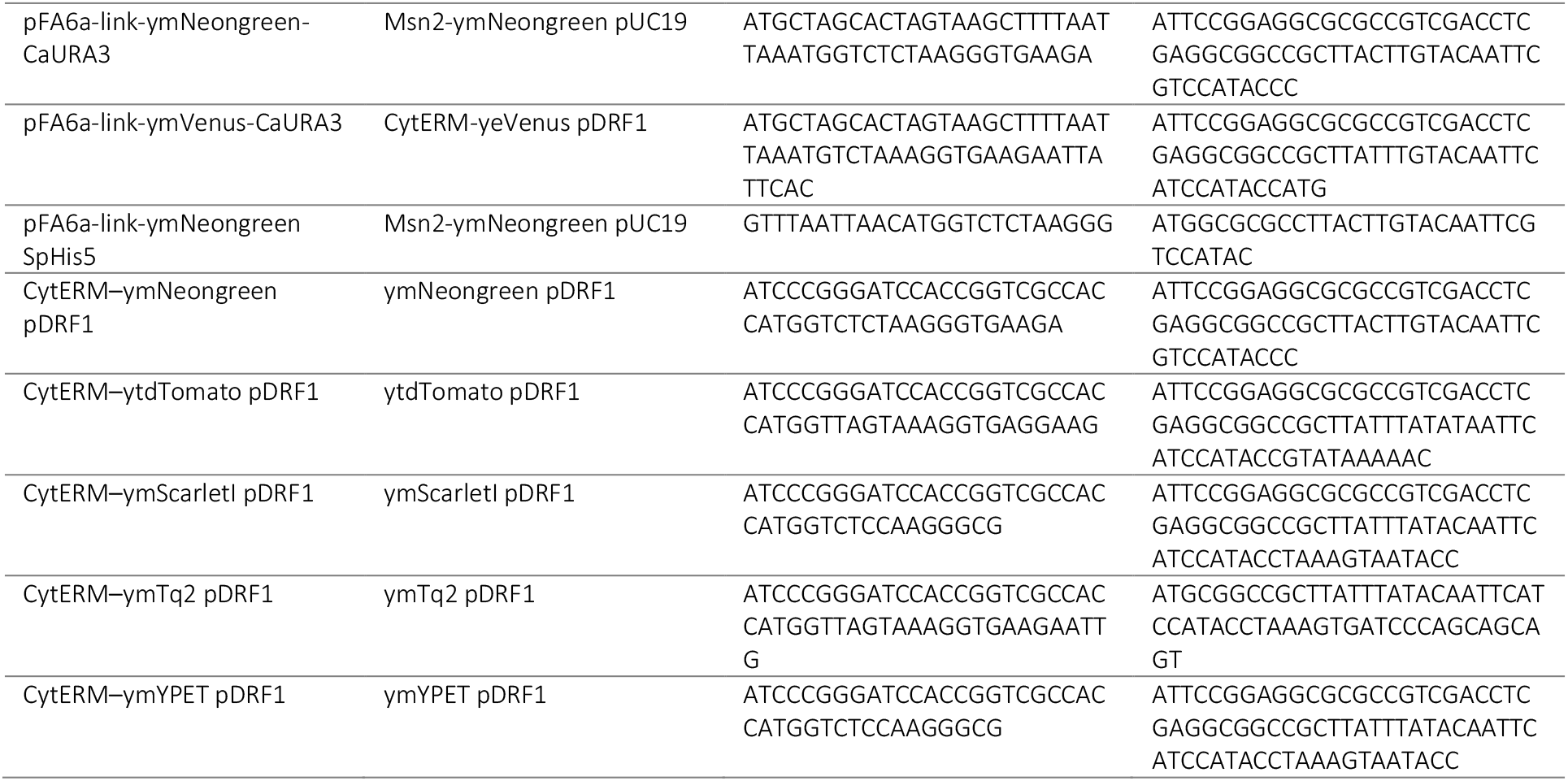
Constructs generated by PCR in this study.

To create yoeCFP-T2A-mCherry, yotagRFPT-T2A-mTq2, yomCherry-T2A-mTq2, yeVenus-T2A-mTq2 and mTq2-T2A-mCherry, PCRs with KOD polymerase were performed according to table 1, the products were digested with NheI and Kpn2I and ligated with T4 ligase into a T2A-mCherry or T2A-mTq2 pDRF1 vector in which the FP N-terminally of T2A was removed by digestion with the same enzymes. This generated yoeCFP-T2A-mCherry, yotagRFPT-T2A-mTq2, yomCherry-T2A-mTq2, yeVenus-T2A-mTq2 and mTq2-T2A-mCherry in pDRF1.

Lastly, eCFP-C1 and mTFP-C1 were digested with NheI and Kpn2I and ligated into mNeongreen-T2A-mCherry in pDRF1 digested with NheI and Kpn2I which replaced mNeongreen with either eCFP or mTFP.

### yFPs

tdTomato, mScarletI, and mYPET (YPET A206K, F208S, E232L, N235D) were codon-optimized and synthesised (Baseclear B.V., Leiden, The Netherlands), generating ytdTomato, ymScarletI and ymYPET. These constructs were digested with NheI and Kpn2I and ligated using T4 ligase into either T2A-mTq2 or T2A-mCherry in which the FP N-terminally of T2A was removed by digestion with the same enzymes. This generated ytdTomato-T2A-mTq2, ymScarletI-T2A-mTq2, ymNeongreen-T2A-mTq2, ymNeongreen-T2A-mCherry and ymYPET-T2A-mTq2 in pDRF1.

Msn2-ymNeongreen and ymTq2Δ9 pUC19 plasmids were codon-optimized and synthesised (Baseclear). A PCR was performed using these constructs according to table 1. Next, the products were digested using NheI and Kpn2I and ligated using T4 ligase into T2A-mTq2 and T2A-mCherry pDRF1 plasmids in which the FP N-terminally of T2A was removed by digestion with the same enzymes, which generated ymTq2-T2A-mCherry, ymNeongreen-T2A-mTq2 and ymNeongreen-T2A-mCherry.

pDRF1 plasmids containing the single yFPs were generated by performing a PCR according to table 1 on Msn2-ymNeongreen, ymTq2Δ9, ymYPET, ytdTomato, ymScarletI in pUC19 plasmids which added a stopcodon at the C-termini. Subsequently, the PCR products were digested with NheI and NotI and ligated with T4 ligase in an empty pDRF1 vector digested with NheI and NotI which generated ymYPET, ymTq2, ymScarletI, ytdTomato and ymNeongreen in pDRF1.

CytERM-ymVenus was created by a mutagenesis PCR according to table 1. Afterwards, pDRF1 containing ymVenus-T2A-mTq2 was constructed by performing a PCR on CytERM-ymVenus according to table 1. Next, the product was digested using NheI and Kpn2I and ligated into a T2A-mTq2 pDRF1 vector in which the FP N-terminally of T2A was removed by digestion with the same enzymes.

pFA6a-link-yFP-CaURA3 plasmids containing the yFPs were generated by performing a PCR according to table 1. Next, the products were digested using PacI and AscI (New England Biolabs) and ligated with T4 ligase into the plasmid pFA6a-link-yomCherry-CaURA3 also digested with PacI and AscI to replace yomCherry with the yFP, which generated pFA6a-link-yFP-CaURA3 plasmids.

pFA6a-link-ymNeongreen-SpHis5 was generated by performing a PCR on Msn2-ymNeongreen pUC19 according to table 1. Next, the product was digested using PacI and AscI and ligated into pFA6a-link-yomKate2-SpHis5 also digested with PacI and AscI (New England Biolabs), replacing yomKate2 with ymNeongreen.

### CytERM constructs

CytERM-dTomato (addgene plasmid #98834) and CytERM-mTq2 (addgene plasmid #98833) were digested using NheI and NotI and ligated into an empty pDRF1 vector digested with the same enzymes which generated CytERM-dTomato and CytERM-mTq2 in pDRF1. CytERM-yeVenus, CytERM-ymNeongreen, CytERM-ytdTomato, CytERM-ymScarletI, CytERM-ymTq2 and CytERM-ymYPET pDRF1 were created by performing a PCR according to table 1. Afterwards, products were digested using XmaI (New England Biolabs) and NotI and ligated with T4 ligase into a CytERM pDRF1 plasmid in which the FP C-terminally of CytERM was removed by XmaI and NotI which generated the CytERM-yFPs.

### Yeast transformation

W303-1A WT W303-1A (MATa, leu2-3/112, ura3-1, trp1-1, his3-11/15, ade2-1, can1-100) yeast cells were transformed according to Gietz and Schiestl, 2007 (Gietz and Schiestl, 2007).

### Characterisation of brightness, day-to-day variation and expression

W303-1A yeast cells expressing the FP-T2A-FP constructs were grown overnight at 200 rpm and 30°C in YNB medium (Sigma Aldrich, Stl. Louis, MO, USA), containing 100 mM glucose (Boom BV, Meppel, Netherlands), 20 mg/L adenine hemisulfate (Sigma-Aldrich), 20 mg/L L-tryptophan (Sigma-Aldrich), 20 mg/L L-histidine (Sigma Aldrich) and 60 mg/L L-leucine (SERVA Electrophoresis GmbH, Heidelberg, Germany). Next, cells were diluted and grown again overnight to mid-log (OD_600_ 0.5-2). Subsequently, samples were put on a glass slide and visualized using a Nikon Ti-eclipse widefield fluorescence microscope (Nikon, Minato, Tokio, Japan) equipped with an Andor Zyla 5.5 sCMOS Camera (Andor, Belfast, Northern Ireland) and a SOLA 6-LCR-SB power source (Lumencor, Beaverton, OR, USA).

CFPs were visualized with a 438/32 nm excitation filter, a 483/32 nm emission filter and a 458 nm long-pass (LP) dichroic mirror. GFPs were visualized using a 480/40 nm excitation filter, 535/50 nm emission filter and a 505 nm LP dichroic mirror. YFPs were visualized with a 500/24 nm excitation filter, a 542/27 nm emission filter and a 520 nm LP dichroic mirror. RFPs were visualized with a 560/20 nm filter, a 610 nm LP emission filter and a 600 nm LP dichroic mirror (all filters from Semrock, Lake Forest, IL, USA). Images were taken using a 60x plan Apo objective (numerical aperture 0.95), 10–30% light power, 2×2 binning and 20–200 msec exposure time at 30°C, dependent on the FP expression. Per FP, fluorescence of 3 biological replicates was recorded. Images were analyzed with FiJi (NIH, Bethesda, MD, USA) and an in-house macro which performs background correction, identifies cells using the Weka segmentation plugin (Arganda-Carreras et al., 2017) and measures the mean brightness of every cell per channel. Data was analysed and visualised using R (R Foundation for Statistical Computing, Vienna, Austria).

### Bleaching kinetics

Cells expressing the FP-T2A-FP constructs were grown and prepared as described for brightness characterisation. The same microscope and filter setups were used as described for brightness characterisation. Bleaching was performed by visualizing the cells every 500 msec, using a 60x plan Apo objective (numerical aperture 0.95), 10% light power, 2×2 binning and 200 msec exposure time at 30°C for 181 frames. Per FP, at least 2 independent bleaching curves were obtained. Afterwards, images were segmented as previously described and the photostability was calculated by dividing the fluorescence of the last time point (frame 181) by the fluorescence of the first time point. Besides, half times were determined by fitting the bleaching curves with a one-phase (equation 1) or two-phase exponential (equation 2) decay formula, with a en b being offsets components for the first and second bleaching component, respectively. X is the time in milliseconds and r and s are the decay rates for each component. Per fit, Bayesian information criterion (BIC) values were obtained for both fits to determine whether the decay curves were mono- or biexponential.

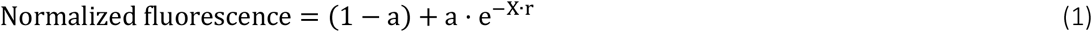

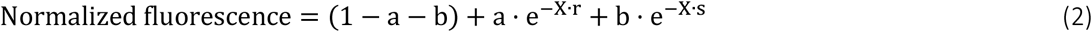

For photochromicity characterisation, cells expressing the FP-T2A-FP were bleached by alternating excitation with the correct wavelength for the FP of interest and a second wavelength (filter setups described in characterisation of brightness), starting with the wavelength for the FP of interest. Bleaching was performed by exposing the cells every 3 sec for both excitation wavelengths, using a 60x plan Apo objective (numerical aperture 0.95), 10% light power, using 2×2 binning and 200 msec exposure time at 30°C. Cells were segmented as previously described and bleaching curves were normalized to the first frame. Model fitting and analysis of photochromism are explained in the supplementary information. Photochromic FPs were selected when an FP had a photochromism value above 50 (photochromism values for each excitation wavelength are shown in table S1).

### pH curves (*in vivo*)

W303-1A yeast cells expressing the FP-T2A-FP constructs were grown overnight at 200 rpm and 30°C in YNB medium containing 100 mM glucose, 20 mg/L adenine hemisulfate, 20 mg/L L-tryptophan, 20 mg/L L-histidine and 60 mg/L L-leucine. Also, W303-1A WT cells were grown overnight at 200 rpm and 30°C in the same medium with 20 mg/L uracil (Honeywell Fluka, Morris Plains, NJ, United States) added. Next, cells were diluted and grown again overnight to an OD_600_ of 1.5-3. Cells were washed twice with sterile water and concentrated to an OD_600_ of 15. Next, cells were diluted 10 times in a 96 wells plate, containing a citrate phosphate buffer (0.1 M citric acid (Sigma Aldrich), 0.2 M-Na_2_HPO_4_ (Sigma Aldrich) set to pH values ranging from 3–8 with 2 mM of the ionophore 2,4-Dinitrophenol (DNP, Sigma Aldrich). Per FP, 3 replicates were used. Cells were incubated for 2 hours to ensure pH equilibration. Next, the fluorescence intensity was measured using a FLUOstar Omega plate reader (BMG labtech, Ortenberg, Germany) using 25 flashes per well at 30°C. Fluorescence of CFPs were obtained by using a 430/10 nm excitation filter and 480/10 nm emission filter, GFPs and YFPs were detected using a 485/12 nm filter and a 520/10 nm emission filter and RFPs were detected using a 544/10 nm excitation filter and a 590 nm long-pass emission filter (all filters from BMG labtech). Cells were corrected for auto fluorescence and normalized to the pH value giving the highest fluorescence. Per FP, a Hill fit (equation 3) was performed to obtain the pKa, the Hill coefficient (steepness of the curve), the offset (the plateau a curve can approach at low pH levels), and the pH value giving an absolute 50% decrease in fluorescence.

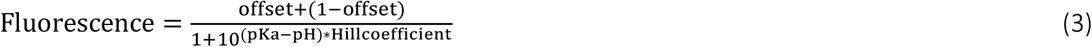

### pH curves (*in vitro*)

50 mL of *E. coli* bacteria expressing mTq2 were grown in LB medium overnight at 200 rpm and 37°C. Next, cells were incubated for 6 hours at 21°C for FP maturation. Cells were harvested by centrifuging at 3220 g for 30 minutes in a swing-out centrifuge using 50 mL tubes. Subsequently, cells were resuspended in 20 mL ST buffer (20 mM Tris, 200 mM NaCl, pH=8) centrifuged again and resuspended in 2 mL ST buffer. The sample was put on ice, lysozyme (1 mg/ml, Sigma-Aldrich) and benzoase nuclease (5 unit/ml, Merck-Millipore) were added and the mixture was incubated for at least 30 minutes on ice. Bacteria were sonicated for 5 mins at 40W and the lysate was centrifuged for 30 mins at 40,000 g and 4 °C. Next, the supernatant was transferred to Ni_2_^+^-loaded His-Bind resin (Novagen (Merck)) and incubated for at least 1 h at 4 °C. The resin was washed three times with 14 mL ST buffer and the FP was obtained by adding 0.5 mL ST buffer containing 0.6 M imidazole. Lastly, the FP solution was dialyzed overnight in 10 mM Tris–HCI pH 8.0 using 3.5 kD membrane tubing (Spectrum Laboratories (Repligen), Waltham, MA, USA). Proteins were snap-frozen and stored at −80 °C.

For pH curves, purified mTq2 protein was thawed on ice. The protein was diluted 10 times in a black μ-clear 96 wells plate (Greiner Bio-One International GmbH, Kremsmünster, Austria) in 100 mM citric acid–sodium citrate buffer (pH 3.0–5.0) or 100 mM phosphate buffer (pH 6.0–8.0) buffer to a final volume of 200 μL. Fluorescence was measured by exciting mTq2 at 430/20 nm and measuring fluorescence at 484/40 nm using a BIO-TEK FL600 Fluorescence plate reader (Biotek, Winooski, VT, USA) at room temperature. Analyses was performed as described for *in vivo* pH characterisation.

### yFP spectra

W303-1A cells expressing the yFPs were grown for at least 2 weeks on 2% agarose plates containing 6.8 gr/L YNB, 100 mM glucose, 20 mg/L adenine hemisulfate, 20 mg/L L-tryptophan, 20 mg/L L-histidine and 60 mg/L L-leucine. An additional set of W303-1A cells without any construct was grown on the the same plates supplemented with uracil. Next, cells were resuspended in selective growth medium (6.8 gr/L YNB, 100 mM glucose, 20 mg/L adenine hemisulfate, 20 mg/L L-tryptophan, 20 mg/L L-histidine and 60 mg/L L-leucine) to an OD_600_ of 3. Subsequently, cells were transferred to a black 96 wells plate (Greiner Bio-One) using 150 μL per well. Per FP, 5 replicates were used. Emission and excitation spectra were recorded using a 1 nm stepsize. For emission spectra, an excitation bandwidth of 16 nm and an emission bandwidth of 10 nm was chosen. For excitation spectra, an excitation bandwidth of 10 nm and an emission bandwidth of 16 nm was chosen. For ymTq2, excitation spectra were recorded from 320 to 530 nm at 565 nm emission, emission was recorded from 428 to 740 nm with excitation set at 398 nm. For ymNeongreen, excitation spectra were recorded from 320 to 540 nm at 570 nm emission, emission was recorded from 480 to 740 nm with excitation set at 450 nm. For ymVenus and ymYPET, excitation spectra were recorded from 320 to 570 with 605 nm emission, emission was recorded from 495 to 740 nm with excitation set at 464 nm For ytdTomato, excitation spectra were recorded from 320 to 626 nm with 662 nm emission, emission was recorded from 530 to 740 nm with excitation set at 500 nm. For ymScarletI, excitation spectra were recorded from 320 to 635 nm with 670 nm emission, emission was recorded from 530 to 740 nm with excitation set at 500 nm. Spectra were corrected for autofluorescence and were normalized to their highest values.

### Fluorescence lifetimes

Cells were grown on 2% agarose plates as described for the yFP spectra. Frequency domain FLIM was essentially performed as described before (Mastop et al., 2017). Briefly, 18 phase images were acquired using a RF-modulated image intensifier (Lambert Instruments II18MD, Groningen, The Netherlands) set at a frequence of 75.1 MHz coupled to a CCD camera (Photometrics HQ, Tucson, AZ, USA) as detector. For cyan FPs, a directly modulated 442 nm laser diode (PicoQuant, Berlin, Germany) was used and for green and yellow FPs a 488 nm argon laser was passed through a RF-modulated AOM (set at 75.1 MHz) for intensity modulation. Emission was passed through a BP480/40 nm filter for cyan FPs and a BP545/30 nm filter for green/yellow FPs. The lifetimes were calculated based on the phase shift of the emitted light (τϕ).

### Oligomerisation tendency

Cells expressing CytERM-dtomato, CytERM-yeVenus, CytERM-ymTq2, CytERM-ymNeongreen, CytERM-ymYPET, CytERM-ymVenus, CytERM-ytdTomato and CytERM-ymScarletI were grown as described for brightness analysis. Next, cells were incubated for at least 1 hour at room temperature, put on a glass slide and visualized using the same setup as described for brightness characterisation. Z-stacks of multiple positions were taken using a Plan Apo λ 100× Oil Ph3 objective (numerical aperture 1.45), 20% light power, 2×2 binning and 100 msec exposure time at 30°C. Per FP, 2 biological replicates were recorded. Images were analyzed with FiJi (NIH, Bethesda, MD, USA) and an in-house macro that performs background correction, makes a Z-projection, identifies cells and OSER structures using the Weka segmentation plugin (Arganda-Carreras et al., 2017) and measures the amount of identified OSER structures per cell. Data was analysed and visualised using R.

### FBPase flow cytometry

According to Gardner and Jaspersen, 2014, a PCR on pFA6a-link-ymNeongreen SpHis5, pFA6a-link-yoEGFP-CaURA3 and pFA6a-link-yoSuperfolderGFP-CaURA3 was performed using KOD Hotstart polymerase with the FW primer CAAATCTTCTATTTGGTTGGGTTCTTCAGGTGAAATTGACAAATTTTTAGACCATATTGGCAAGTCACAGGGTGACGGTGCTGGTTTA and the RV primer ATACAGATTTTTTTTTTCGCGTACTAAAGTACAGAACAAAGAAAATAAGAAAAGAAGGCGATCATTGAATCGATGAATTCGAGCTCG. Next, the products were transformed in CEN.PK2-1C WT (MATa; ura3-52; his3-Δ1; leu2-3,112; trp1-289; MAL2-8c SUC2, obtained from Euroscarf), generating CEN.PK2-1C + FBP1-yoeGFP (MATa; ura3-52; his3-Δ1; leu2-3,112; trp1-289; MAL2-8c SUC2 FBP1-yoeGFP (URA)), CEN.PK2-1C + FBP1-yosfGFP (MATa; ura3-52; his3-Δ1; leu2-3,112; trp1-289; MAL2-8c SUC2; FBP1-yosfGFP (URA)) and CEN.PK2-1C + FBP1-ymNeongreen (MATa; ura3-52; his3-Δ1; leu2-3,112; trp1-289; MAL2-8c SUC2; FBP1-ymNeongreen (HIS)). CEN.PK2-1C + FBP1-yoeGFP and CEN.PK2-1C + FBP1-yosfGFP were grown overnight at 30°C and 200rpm in 1× YNB medium containing 20 mg/L L-histidine, 60 mg/L L-leucine, 20 mg/L L-tryptophan and 50 mM phthalate buffer at pH 5 (adjusted with KOH) and 100mM glucose. CEN.PK2-1C + FBP1-ymNeongreen was grown overnight at 30°C and 200rpm in 1× YNB medium containing 20 mg/L L-uracil, 60 mg/L L-leucine, 20 mg/L L-tryptophan and 50 mM phthalate buffer at pH 5 (adjusted with KOH) and 100 mM glucose. CEN.PK2-1C WT was grown overnight at 30°C and 200rpm in 1× YNB medium containing 20 mg/L L-histidine, 20 mg/L L-uracil, 60 mg/L L-leucine, 20 mg/L L-tryptophan and 50 mM phthalate buffer at pH 5 (adjusted with KOH) and 100 mM glucose. Next, cells were diluted in medium containing 2 mM glucose and 100 mM ethanol or medium containing 100 mM glucose and grown again overnight to an OD_600_ of 1. Next, GFP was measured using a BD Accuri C6 Flow Cytometer (Becton, Dickinson and Company, Franklin Lakes, NJ, USA). Per sample, 50 μL was run on medium flowrate with a maximum of 10.000 events per second. The threshold was set on a forward scatter height (FSC-H) of 80.000 and fluorescence was recorded using 488 nm excitation and an emission filter of 533/30 nm. Data was analysed and visualised using R.

## Results

### *In vivo* brightness and photostability

The most important criterium for choosing an FP is its brightness, as this largely determines the fluorescent signal that can be obtained. To obtain the practical brightness, we linked 27 of the mostly used FPs to either mTurquoise2 (mTq2) or mCherry with a viral T2A peptide, as was previously done for mammalian cells (Goedhart et al., 2010). We did not include the yeast-optimized GFP Envy as it is known to be dimeric (Bajar et al., 2016; Slubowski et al., 2015). The linkage through the T2A peptide ensures the equimolar expression of the two FPs (Goedhart et al., 2011) required for quantitative comparisons. Normalizing the FP of interest to the expression levels of mTq2 or mCherry (the control FPs) gives the practical brightness of an FP in yeast (Figs 1 and 2, Table S1).

**Figure 1.**
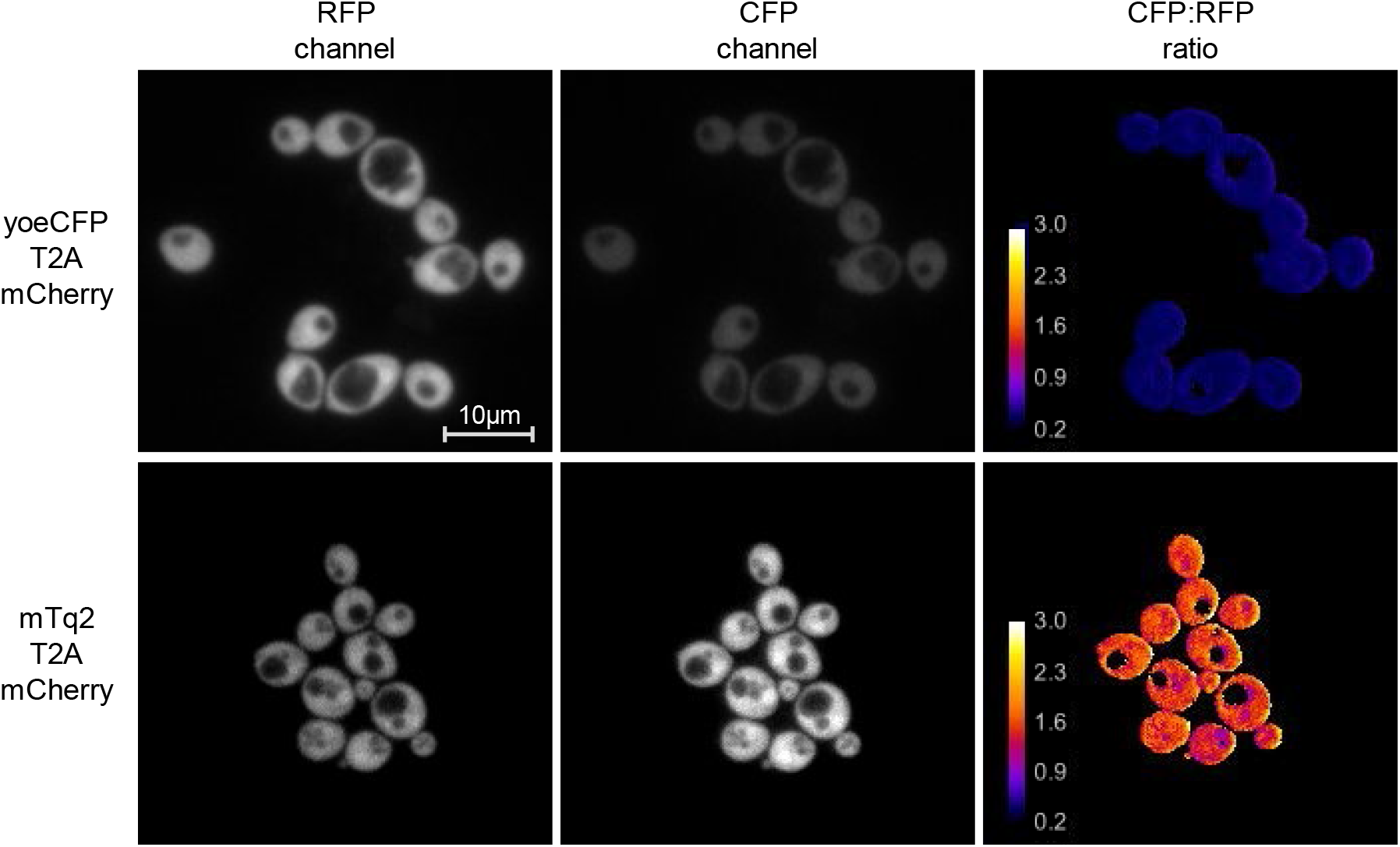
Example of practical brightness quantification using the T2A peptide linker. Cells expressing either yoeCFP-T2A-mCherry or mTq2-T2A-mCherry were grown to midlog and visualized using a widefield microscope. yoeCFP shows a low brightness compared to mCherry. In contrast, mTq2 shows a higher brightness than mCherry. Calibration bar indicates the ratio value when dividing the CFP by the RFP channel (i.e. the relative brightness to mCherry).

**Figure 2.**
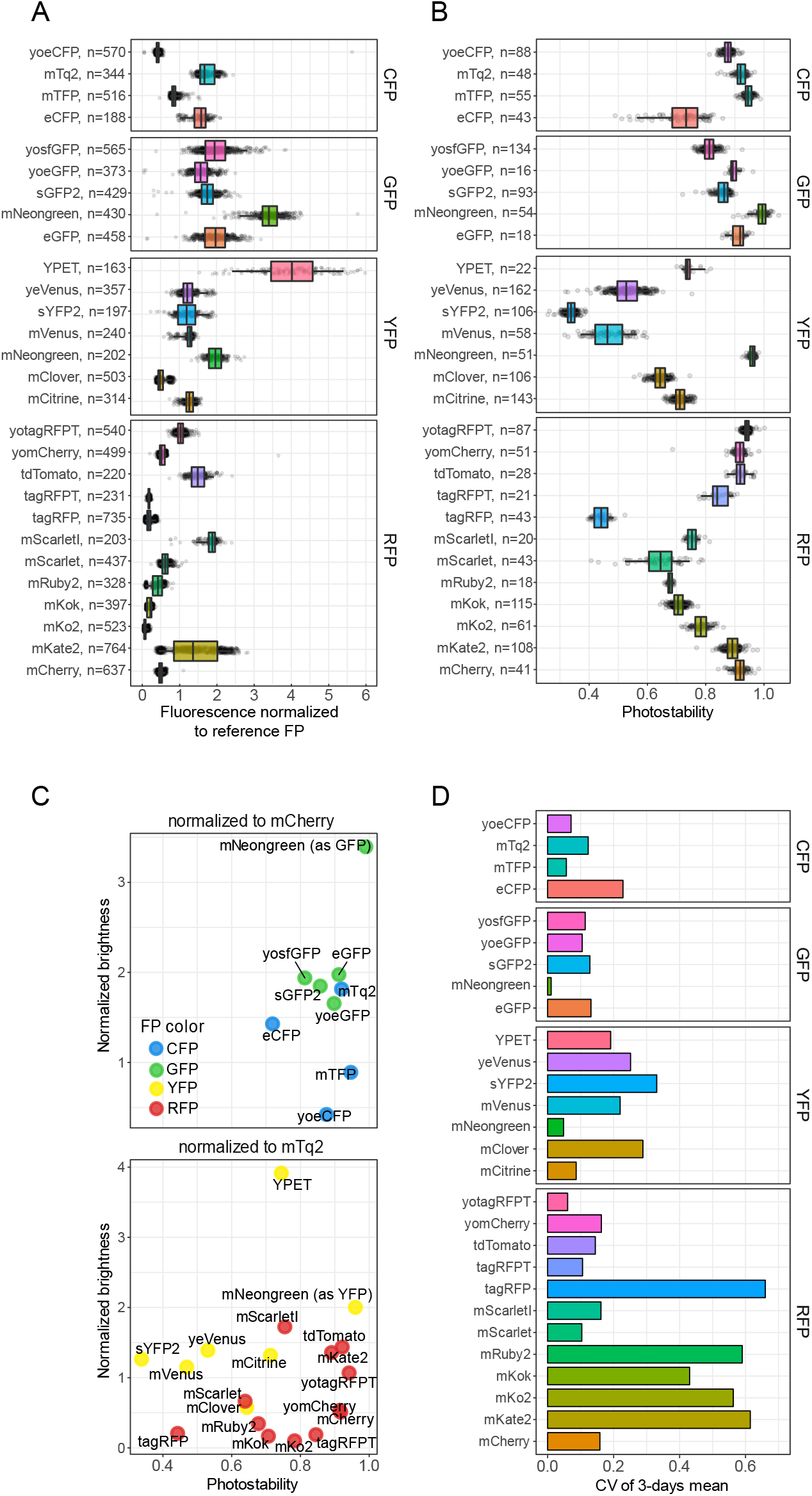
In vivo brightness and photostability of FPs. A) In vivo brightness of FPs measured by normalizing the fluorescence intensity of single-cells expressing FP-T2A-FP and dividing the fluorescence of the FP of interest to the control FP (either mTq2 or mCherry). B) Photostability of FPs. Per FP, a time-lapse movie was recorded and the photostability was measured as the fluorescent fraction of the last time frame compared to the first frame. Dots represent relative brightness or photostability of an individual cell, boxes indicate median with quartiles, whiskers indicate the 0.05–0.95 fraction of the datapoints. C) Overview of the brightness and photostability of all characterised FPs. D) Coefficient of variation (CV) of the mean brightness of 3 days as an indication of day-to-day variation.

As can be inferred from figure 2A and 2C, the differences in practical brightness between spectrally similar fluorescent proteins were substantial. A relatively low brightness was observed for mTFP, mClover, tagRFP, mScarlet, mRuby2, mCitrine and mVenus. In contrast, eCFP, mTq2, mNeonGreen, YPET, mScarlet-I and mKate2 showed a relatively high practical brightness. In addition, tdTomato also showed high brightness. However, tdTomato is known to mature badly in mammalian cells in which it shows a large fraction of unmature green fluorescent proteins (Shaner et al., 2004; Van der Krogt et al., 2008). Yet, this did not occur in yeast and tdTomato is therefore a useful red FP in yeast. The altered brightness of yoeCFP and yotagRFP-T compared to their non-codon optimized variants might be due to these constructs not being completely identical. We also measured day-to-day variation, depicted by the coefficient of variation (CV) of the mean of each day. We found various YFPs and RFPs to have a large day-to-day variation (Fig 2D). These FPs will give broader distributions or different readouts when used at different days due to intrinsic brightness variations between days.

Next to brightness, the photostability of FPs is often considered when choosing an FP as this determines how long an FP can be visualized. To assess photostability, we imaged cells expressing the FPs using low amounts of widefield exposure as this resembles real experiments best. All fluorescence was normalized to the first frame and the photostability was determined as the fluorescence fraction still present at the last frame of the bleaching experiment (Figs 2B and 2C). Various FPs in the red and yellow spectrum showed a low photostability whereas most of the CFPs and GFPs showed a high photostability. We found that the relative photostability (i.e. compared to other FPs) of mVenus and yosfGFP was lower than previously determined (Cranfill et al., 2016). In contrast, YPET, mCitrine, mKate2 and tdTomato were relatively more photostable than reported *in vitro.* The most photostable FPs are mTFP, mNeongreen, tdTomato, mCherry and yotagRFPT. YPET and mCitrine were the most photostable yellow FPs (besides mNeongreen) but still showed considerable bleaching. Combining FP brightness with photostability resulted in YPET, mNeongreen, mTq2, tdTomato and mScarletI as the best performing FPs.

### Photochromism

Although photobleaching experiments give information about FP photostability, these single-wavelength bleaching kinetics give an incomplete picture of FP behaviour in time. Nowadays, FPs are often used simultaneously so that FPs are excited at various wavelengths. Exposing an FP to multiple wavelengths can induce or accelerate both reversible and irreversible photobleaching (Bindels et al., 2017; Dean et al., 2011; Dickson et al., 1997; Shaner et al., 2008; Sinnecker et al., 2005). Moreover, another photophysical phenomenon of FPs is photoswitching or photoconversion into a spectrally different species (Kremers et al., 2009). The term that we will use for any of these effects is photochromism. We identified photochromic FPs and focussed on the effect of different excitation wavelengths on photochromism.

We assessed photochromic behaviour of all FPs in our library (Figs 3 and S3). We did this by fitting bleaching curves of single-wavelength and dual-wavelength bleaching data to a mathematical model. This model includes 3 FP states: a natural (nat), a reversible dark (dark), and an irreversible dark state (irrdark) (fig 3A). In the natural state, the FPs are fluorescent. FPs can transition from the natural state to the dark state, both by light exposure and spontaneously. In the dark state, the FPs are not fluorescent, but can return to the natural state both spontaneously and by excitation light. For the different wavelengths that can be combined for an FP, we fitted different rate constants for these transitions, as well as for the spontaneous transitions. Lastly, FPs can also transition from the dark state to an irreversible dark state in which the FP stays non-fluorescent. Using this small model, we were able to fit all obtained bleaching kinetics for each FP (Figs 3C, S3). The model can therefore be used to correct for complex bleaching kinetics occurring with photochromic FPs.

**Figure 3.**
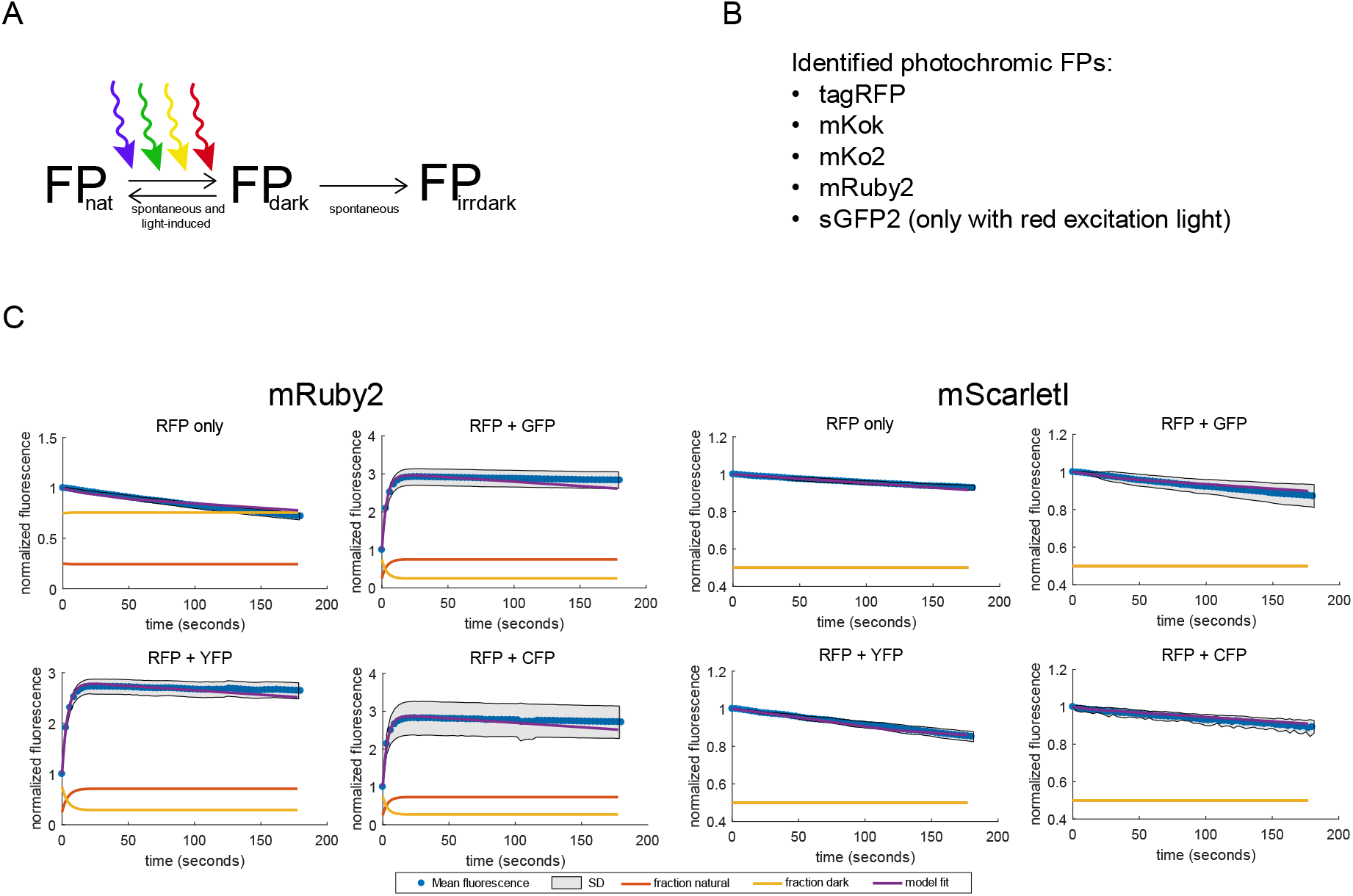
Photochromism characterisation, determined by recording bleaching data of FPs using single-wavelength and dual-wavelength excitation. A) Model used to fit the bleaching data and obtain photochromism parameters. B) The identified photochromic FPs. C) Bleaching plots including model fitting of the photochromic mRuby2 and the non-photochromic mScarletl. Dots represent mean fluorescence values at the specific time point, normalized to the first frame. Shades indicate standard deviation. Red and yellow lines indicate the fitted natural (fluorescent) FP fraction and reversible dark FP fraction, respectively. Blue lines indicate the fitted mean fluorescence, normalized to the first frame. Used wavelengths are shown above each graph.

With the model we were also able to show that tagRFP and mRuby2 are photochromic, which is in agreement with other studies (Fig. 3B) (Bindels et al., 2017; Dean et al., 2011; Shaner et al., 2008). We also identified mKok and mKo2 as photochromic, in agreement with an earlier observation that blue light triggers photoconversion of these FPs (Goedhart et al., 2007; Mastop et al., 2017). Lastly, we observed photochromism of sGFP2. Interestingly, photochromic behaviour of most FPs only occurs when using wavelengths shorter than the optimum wavelength for the FP. This suggests that the transition from the reversible dark state to the natural state requires more energy than the energy required to excite the fluorescent protein. To test the effect of each specific excitation wavelength on photochromism, we systematically determined the photochromic sensitivity of an FP for any of the other wavelengths (i.e. CFP, GFP, YFP or RFP excitation wavelengths) that can be combined with that FP (for example, YFP photochromism is not determined in combination with a GFP excitation wavelength as a YFP and GFP cannot be combined for dual colour experiments). Our data indicates that the specific second wavelength used does not largely affect photochromism effects. The only requirement is a wavelength smaller than the optimum wavelength for the specific FP. The identified photochromic FPs can create biased fluorescence readouts and should be used carefully. When these FPs are used with multiple excitation wavelengths, bleaching corrections can be performed with our supplied model.

### pH stability

Next, we determined the brightnes of all 27 FPs at different pH values to determine which are usable in acidic environments or when intracellular pH is dynamic. We measured pH-induced quenching *in vivo* by incubating cells in a citric-acid phosphate buffer with pH values ranging from 3–8. In order to ensure equilibration of the intracellular pH with the buffer we added the ionophore 2,4-DNP, which is known to remove the pH gradient in yeast (Thevelein et al., 1987). FP fluorescence was measured and a Hill fit was performed to obtain the pKa values and Hill-coefficients (Figs. 4A, S1 and Table S1). We also determined the pH value that gives an absolute 50% decrease compared to the pH value with the highest fluorescence, as the pKa value does not always give the pH value with 50% fluorescence decrease when FPs show offsets (Fig 4C). Examples are sYFP2, mScarletl, mScarlet or mClover, which have different pH_50%_ value compared to their pKa. We also found 7 FPs with a pKa that differed more than 0.5 compared to previously published *in vitro* data (Table S1). mTq2, mTFP and mClover showed increased pH sensitivity; In contrast, eCFP, YPET, mKate2 and mScarletl showed decreased pH sensitivity. We confirmed the difference between our *in vivo* assessment with *in vitro* assessments (Fig. 4B). Therefore, *in vitro* quenching characterisation is not representative for the *in vivo* FP behaviour, at least not in yeast cells.

**Figure 4.**
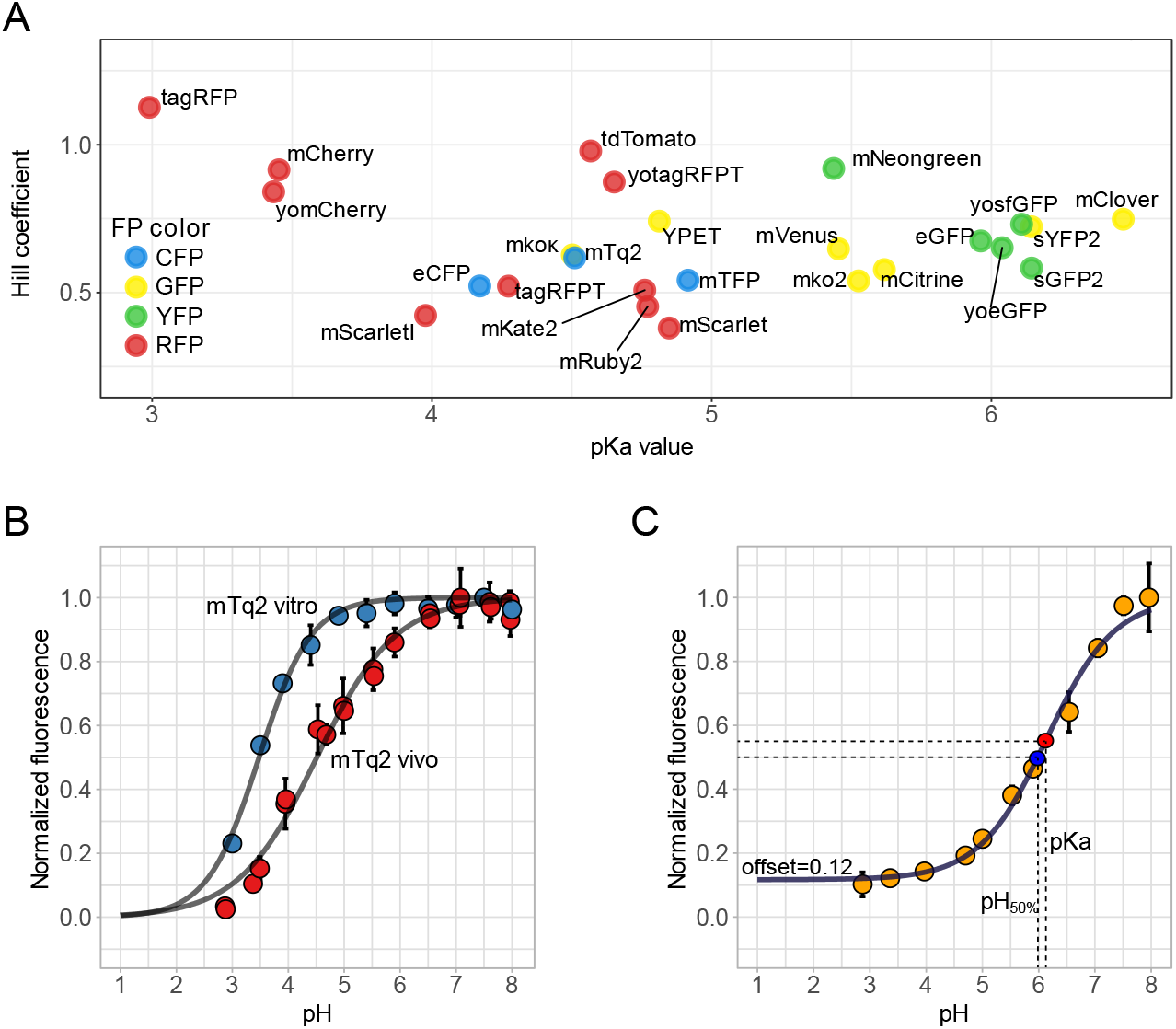
pH sensitivity of FPs. A) Yeast cells were incubated for 2 hours in citric-acid/Na_2_HPO_4_ buffers set at pH 3-8 with 2 mM 2,4-DNP and fluorescence was measured using a fluorescent plate reader. Per FP, at least 3 technical replicates were measured. Afterwards, fluorescence was normalized to the pH giving the highest fluorescence and a Hillfit was performed to determine the hill coefficient and pKa value, plotted at the y- and x-axis, respectively. B) mTq2 is an example of an FP that shows different pH sensitivity. pH calibration in vitro was performed using purified proteins in a Citric Acid – Sodium Citrate buffer (pH 3 – 5.4) and a NaH_2_PO_4_/Na_2_HPO_4_ 0.1 M buffer (pH 5.9-8). Dots represent mean of at least 3 replicates, error bars indicate SD. C) pH curve of sYFP2 which shows an offset (fluorescence plateau) at low pH. This offset gives different values of the pKa (red point, which is the pH that gives a 50% decrease between 1 and the offset) and the pH_50%_ which gives an absolute 50% decrease in fluorescence (blue point). Dots represent mean of at least 3 replicates, error bars indicate SD.

To be insensitive to pH changes, the Hill coefficient of a pH-curve should be high and the pKa should be low; then the curve shows a plateau where the fluorescence does not change at physiological pH values. Fig. 4A shows that CFPs and RFPs show a low pKa whereas most YFPs and GFPs have a higher pKa. Of the ones with a low pKa, only yomCherry, mCherry, YPET, tagRFP, yotagRFPT and tdTomato show also high Hill coefficient. Therefore, these FPs are most pH-insensitive. In conclusion, pH sensitivity of many FPs is different *in vivo* than *in vitro.* Of the identified bright and photostable FPs, tdTomato is the most pH robust. Although less pH robust, mTq2, mNeongreen and YPET are still the best CFP, GFP and YFP variants, respectively.

### Yeast codon optimization improves expression and changes FP characteristics

In our assays we identified mTq2, mNeongreen, YPET, tdTomato and mScarletI as the best performing FPs for yeast. These FPs are bright, photostable, pH robust and have a low day-to-day variation. However, most of these FPs have a low expression (Fig 5C), making them unsuitable for either plasmid-based overexpression or tagging of proteins in yeast. Therefore, we created codon-optimized variants of these FPs, which we named yeast FPs (yFPs). We also included ymVenus as this FP was also an acceptable YFP. Since YPET is known to be not monomeric, we synthesised yeast monomeric YPET (ymYPET: YPET A206K, F208S, E232L, N235D). The novel yFPs were rescreened for photostability, pH stability and brightness (Fig 5). ymVenus, ymNeongreen and ymTq2 showed a significant increase in brightness whereas ytdTomato and ymScarletI had a decreased brightness compared to the non-optimized versions (Tukey HSD, p<0.01). ymYPET appeared 4 times less bright, indicating that the introduced mutations affect its brightness. Still, ymYPET is one of the brightest YFPs available. Photostability remained the same for yFPs, except for ymVenus and ymYPET (Tukey HSD, p>0.01). Fluorescent lifetimes were within a 0.1 ns range with previously reported values. Spectral properties, pH stability and day-to-day variation were comparable as well (Figs. 5E, 7 & S2, table 1 and S1). As anticipated, the expression of yFPs is largely increased (Fig. 5C), showing the importance of optimized codon-usage on protein expression. Lastly, to determine whether the yFPs are truly monomeric, we performed an OSER assay by fusing the yFPs to the first 29 amino acids of cytochrome p450, which is targeted to the endoplasmic reticulum (Costantini et al., 2012). Non-monomeric FPs will form multimeric complexes which generates bright spots in a cell, named whorls. The amount of whorls or percentage of cells with a whorl is used as an estimate for monomerism. Since this assay was developed for mammalian cells, we tested the assay in yeast by including yeVenus and dTomato, which are known to be nonmonomeric (Cranfill et al., 2016; Shaner et al., 2004). As expected, yeVenus and dTomato showed a significant higher amount of whorls per cell compared to all yFPs, except for ymVenus and ytdTomato (Fig. 6). ymTq2, on the other hand, showed only a significantly lower amount of whorls compared to yeVenus (Fig. 6, Tukey HSD, *α* = 0.01). We conclude that only ymVenus (yeVenus A206K) and ytdTomato are not monomeric and are therefore inferior to ymYPET and ymScarletI, respectively. Fig 6 shows that all other codon-optimized FPs are monomeric according to our assay and therefore suitable for use in yeast cells.

In summary, codon-optimization may change the protein properties and should therefore be followed by a comprehensive *in vivo* characterization.

**Figure 5.**
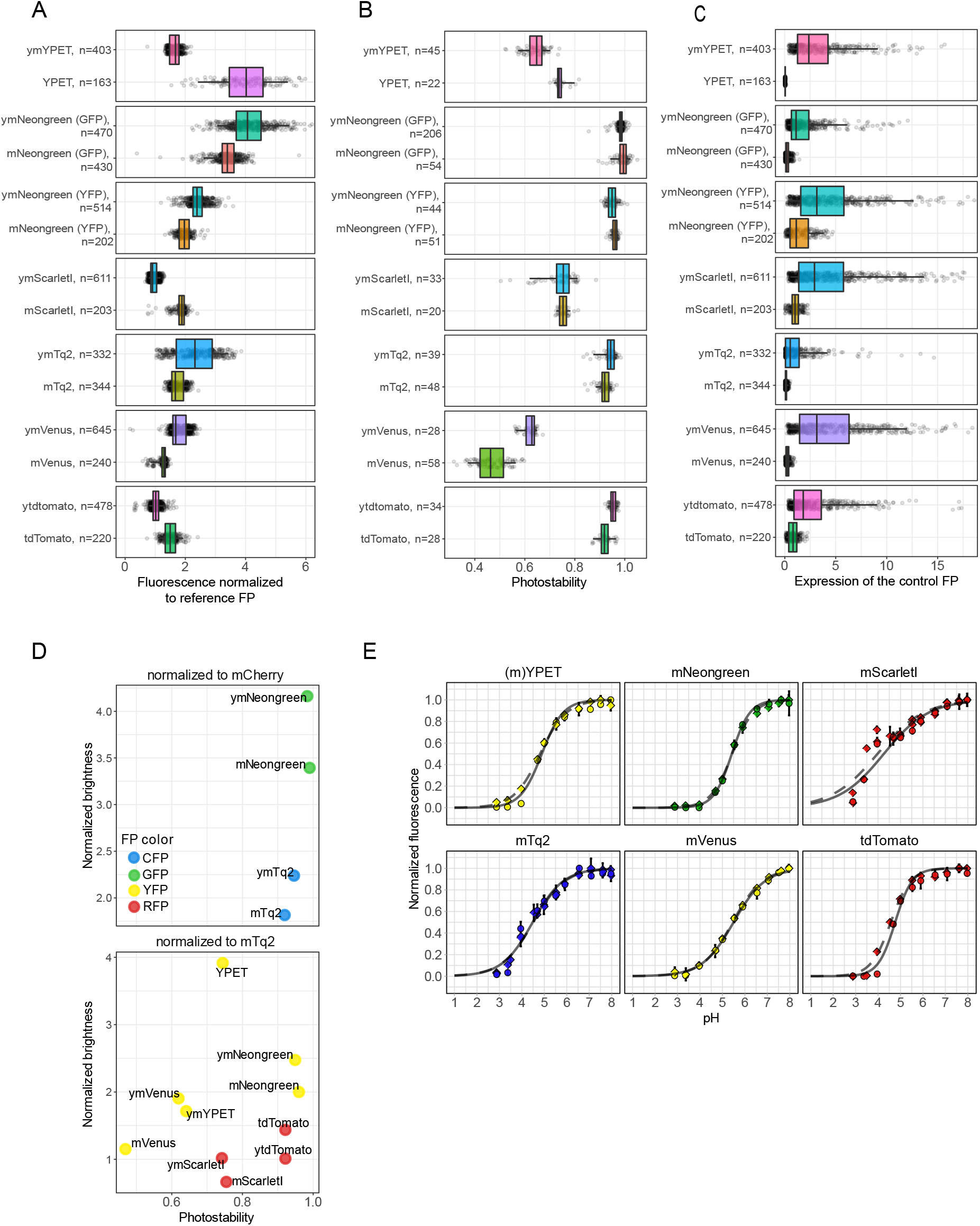
Comparison of yeast codon-optimized FPs (yFPs) versus the conventional FPs. A) Comparison of FP brightness. Dots represent FP brightness of an individual cell, boxes indicate median with quartiles whiskers indicate the 0.05–0.95 fraction of the datapoints. B) Comparison of photostability. Dots represent FP photostability of an individual cell, boxes indicate median with quartiles, whiskers indicate the 0.05–0.95 fraction of the datapoints. C) Comparison of FP expression. Dots represent fluorescence of the reference FP of an individual cell (divided by the total amount of exposure), boxes indicate median with quartiles, whiskers indicate the 0.05–0.95 fraction of the datapoints. D) Overview of both the photostability and brightness of the yFPs and their FP counterparts. E) pH sensitivity of the yFPs compared to the FPs. Dashed lines with circles show yFPs fit and the data points, respectively. Solid lines with rhombic points show FPs fit and the data points, respectively.

**Figure 6.**
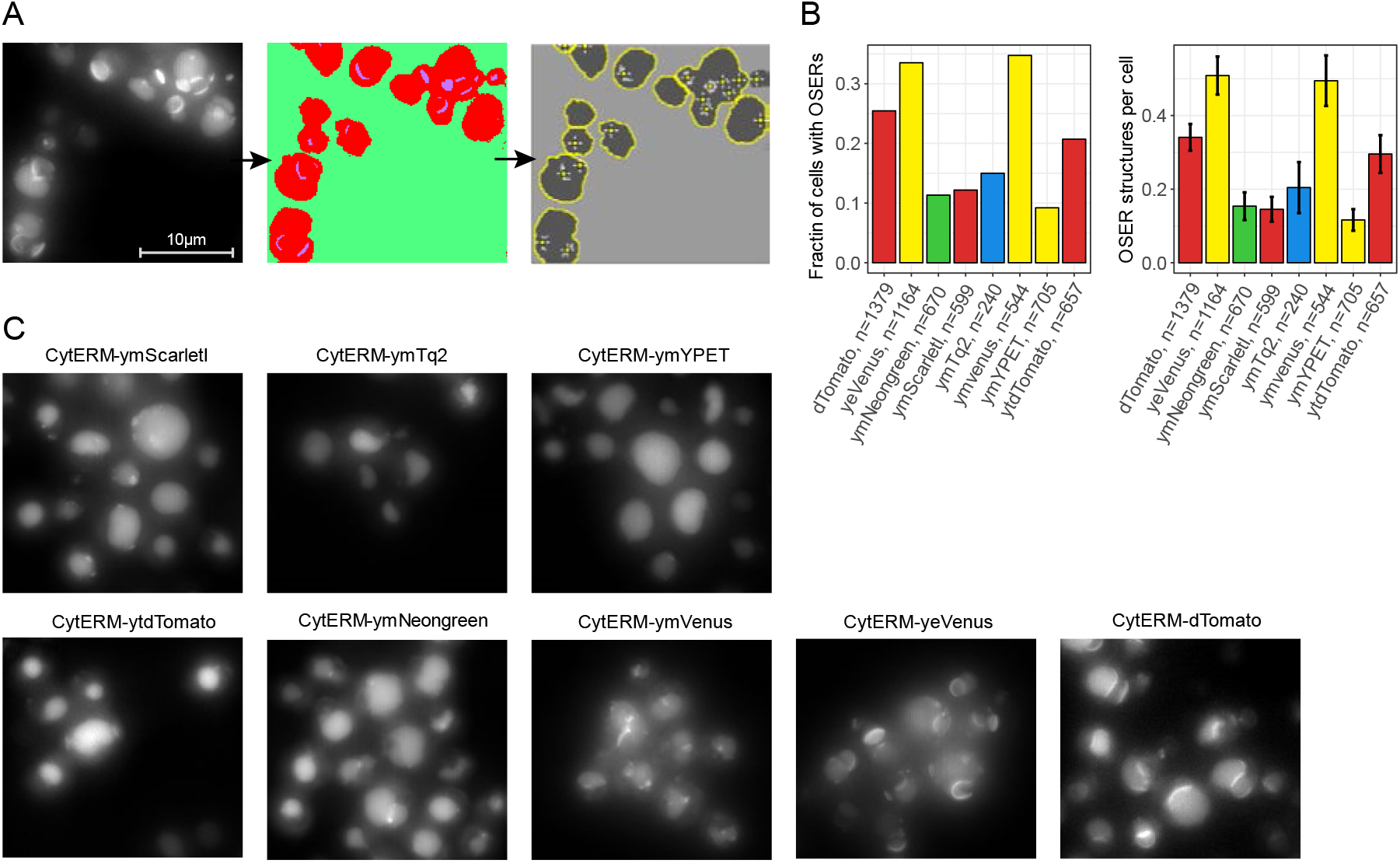
Monomerism of the yFPs. Cells expressing Cyterm-(y)FP were grown overnight at 30 °C and incubated for at least 1 hour at room temperature before microscope visualization. A) Example of cells showing OSER structures and the pipeline to count OSER structures. B) Monomerism of various FPs depicted by the fraction of cells with an OSER structure (left graph) and amount of OSER structures per cell (right graph). Error bars indicate 95CI. C) Examples of the OSER assays per FP (max intensity projection of the Z-stack is shown).

**Figure 7.**
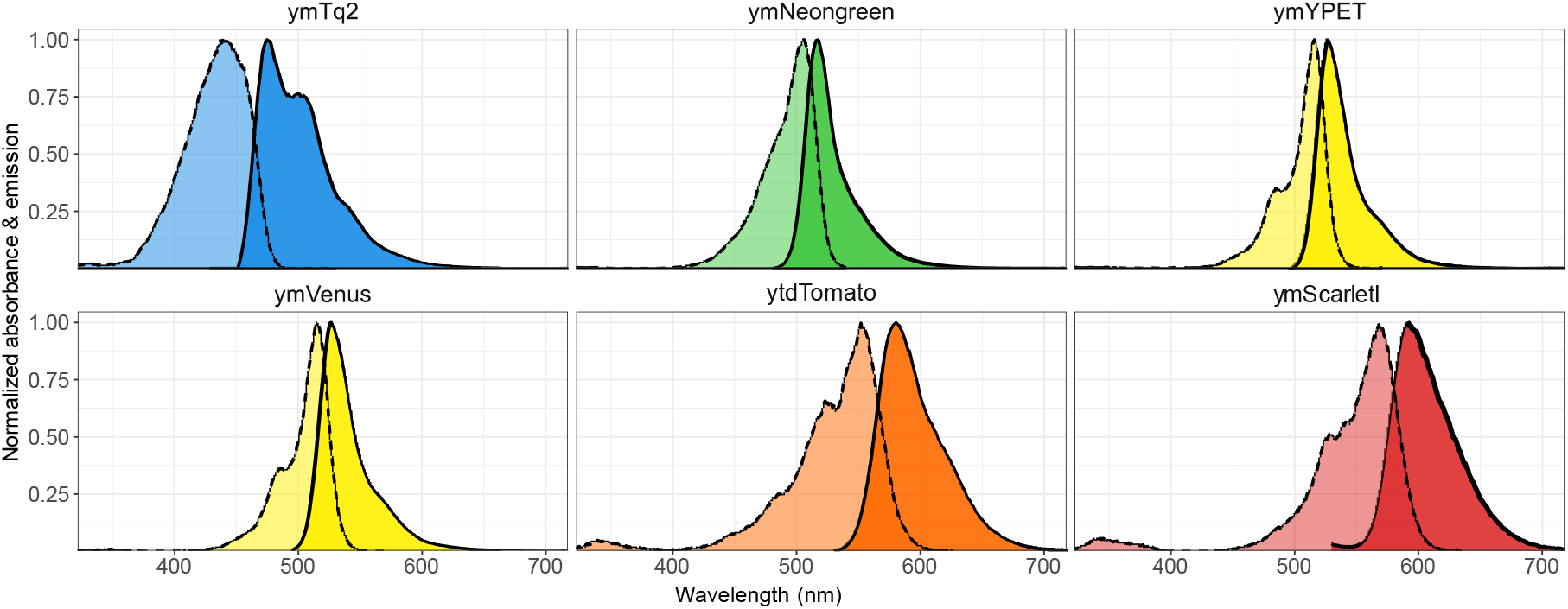
Excitation and emission spectra of the best performing yFPs. Spectra were normalized to the highest excitation or emission value per yFP. Dotted lines show excitation spectra. Solid lines show emission spectra.

**Table 1.**
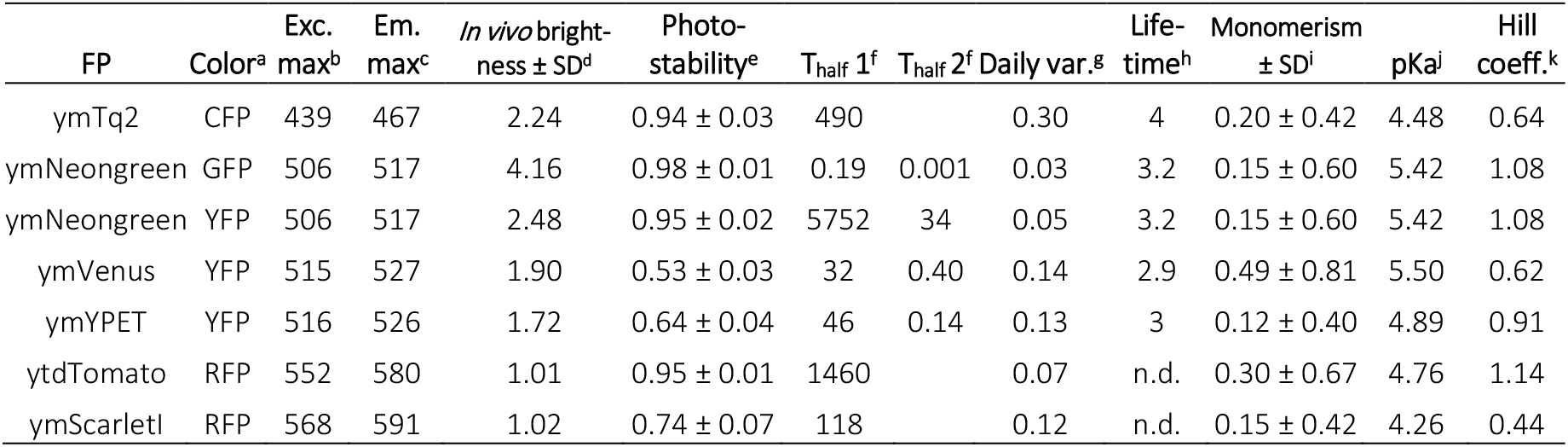
In vivo properties of the yFPs. ^a^Spectral color class. ^b^Excitation maximum. ^c^Emission maximum. ^d^Brightness in budding yeast normalized to either mCherry (for CFPs and GFPs) or mTq2 (for YFPs and RFPs). ^e^Fluorescent fraction remaining at the last timeframe, normalized to the first frame. ^f^T-half times in seconds obtained via a one-phase or two-phase exponential decay fit. ^g^Coefficient of variation (CV) of the mean brightness of 3 days as an indication of day-to-day variation. ^h^Fluorescence lifetime in nanoseconds determined by frequency domain. ^i^Monomerism depicted by the mean amount of OSER structures per cell. ^j^pH value giving 50% decrease in fluorescence. ^k^Hill coefficient of the pH stability. ^l^pH value giving an absolute decrease of 50% in fluorescence. Abbreviations: SD; standard deviation, n.d.; not determined.

### yFPs show improved readout and enable fluorescent-based quantification of the enzyme FBPase

One benefit of using yeast is the opportunity to tag proteins endogenously. To do so, we created a library of tagging vectors containing the yFP palette. We tested whether the newly codon-optimized FPs can improve experimental readouts with one of these tagging vectors. FBPase, an enzyme expressed in yeast under gluconeogenic conditions (e.g. growth on ethanol) was fused with yoeGFP, yosfGFP and ymNeongreen. We compared the green fluorescent signal of these strains when growing on either glucose (without FBPase expression) or ethanol (with FBPase expression) using flow cytometry (Fig. 8). Fig. 8 shows that yosfGFP (43% overlap) and yoeGFP (51% overlap with WT) are inferior to ymNeongreen (1% overlap with WT) regarding GFP signal when fused to FBPase. The signal of ymNeongreen is not due to aspecific or leaky expression as this strain shows the same low (background) fluorescence as the WT strain when grown in glucose. Therefore, the use of the newly developed ymNeongreen enables flow cytometry-based monitoring of FBPase expression in yeast cells, which is not feasible using the conventional FPs. By this, we show that the use of the codon-optimized FPs can greatly improve fluorescent-based readouts.

**Figure 8.**
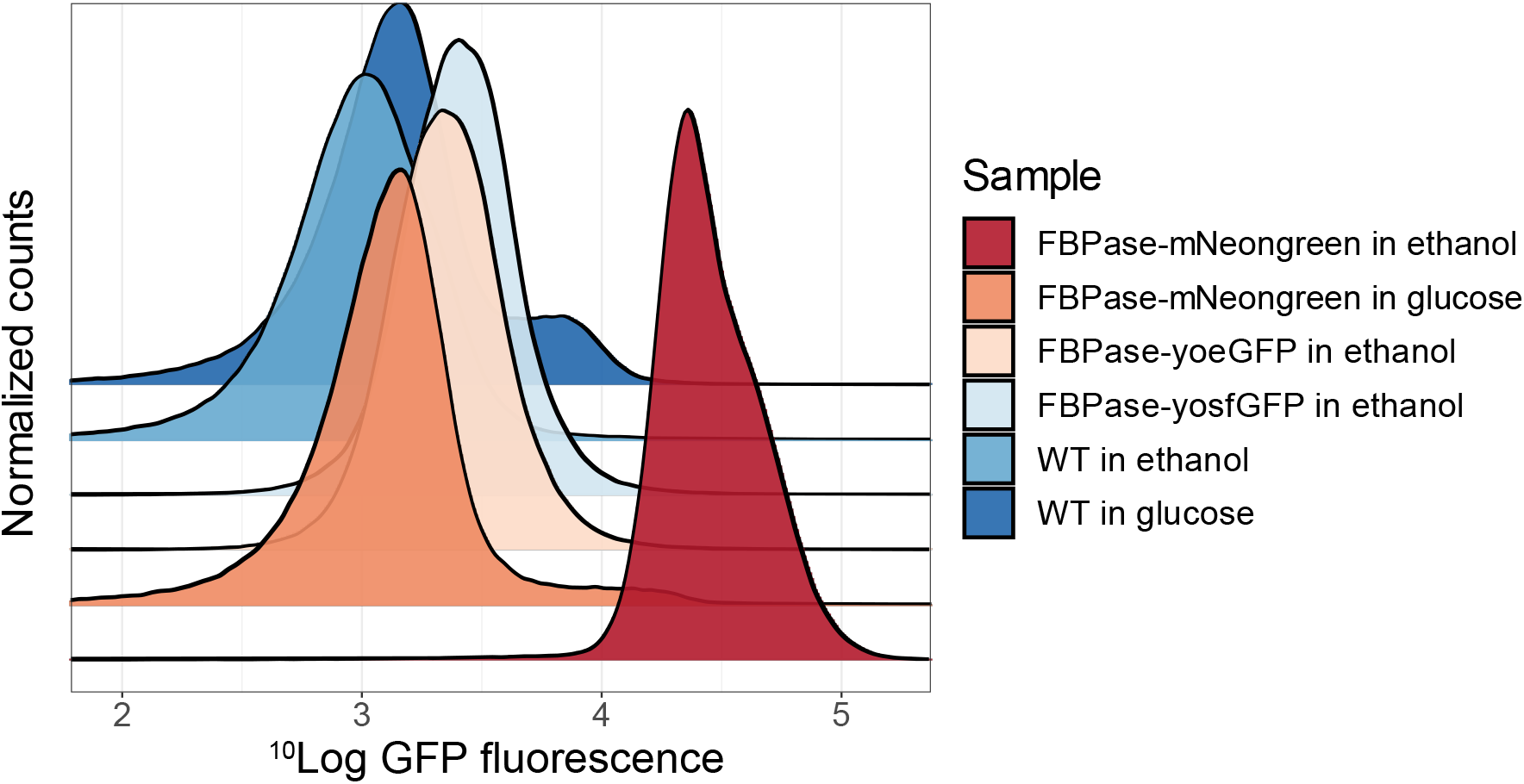
Normalized frequency plots of GFP fluorescence of Cen.PK yeast cells with FBPase-yosfGFP, FBPase-yoeGFP, FBPase-ymNeongreen or WT. Cells were grown in YNB medium containing either glucose (no FBPase expression), or ethanol (with FBPase expression) and GFP fluorescence was measured using flow cytometry.

## Discussion

Comprehensive data on the *in vivo* performance of FPs in eukaryotes grown at modest temperature (30°C) is largely unavailable. Yet, these organisms are widespread in the eukaryotic kingdom. In particular, *Saccharomyces cerevisiae* is often used as model eukaryotic organism and as workhorse in biotechnology. Therefore, we systematically characterised FPs *in vivo* in budding yeast. Key in our approach is the T2A linker peptide, through which we determined relative brightness of FPs by comparing it to the control FP. In the same strains, we tested FPs for photostability and pH sensitivity. Lastly, we provide a comprehensive and systematic analysis of the tendency of FPs for photochromism and we provide a model to correct for it when such FPs are still being used in multicolor experiments.

The brightness of FPs shows large differences for various FPs compared to *in vitro* data (Table S1). These results show that *in vivo* conditions indeed affect FP behaviour. We suspect -but have not tested-that temperature is an important factor, since yeast grows optimally at 30°C, but nearly all FPs are optimized for maturation at 37 degrees (Balleza et al., 2018; Griesbeck et al., 2001; Nagai et al., 2002). Interestingly, all bad performing FPs indeed show a relatively bad maturation at 32 degrees as previously studied (Balleza et al., 2018). Yet, two observations suggest that the effect of temperature on FP performance is not generally applicable. First, mKate2 shows bad maturation at low temperatures but is bright in our analysis. Second, in *C. Elegans,* that also grows at lower temperatures, mNeongreen is less bright than in our study (Heppert et al., 2016). This shows that FPs should be characterised *in vivo* in each organism. Importantly, the practical brightness reported here depends on the excitation and emission filters that were used. Since FPs do not have identical spectra, changing the filters will affect the brightness. On the other hand, the filters that we used are commonly used. In any case, the plasmids that we have constructed allow anyone to rapidly determine the practical brightness under their own specific conditions.

One other property we could assess with our method is the variation of brightness between different days. Various RFPs and YFPs show a high day-to-day variation which make these FPs less desirable for quantification (Fig. 2D). One explanation could be hampered FP maturation due to the low temperature, as most of the FPs with a high variation are also known for bad maturation (Balleza et al., 2018). However, other factors such as translation and folding efficiency (affected by chaperones) could also affect maturation efficiency, which increases day-to-day variation. We recommend to carefully select FPs with a low high day-to-day variation when the study requires accurate quantitative data on noise levels (protein expression distribution) or on the actual protein levels.

We found that most FPs show comparable photostability as previously characterised. Importantly, FPs with a high photostability (e.g. ymNeongreen) give meaningless T_half_ times, which is due to the lack of bleaching. Therefore, we believe that the photostability parameter given is a better approximation for photostability than the obtained T_half_ times. Photostability is also largely dependent on the setup and specific exposure type and is therefore difficult to compare with previously obtained data (Cranfill et al., 2016). Therefore, it is recommended that users do a brief characterisation with their own setup. Still, our photostability data gives an indication of the photostability of the FPs. A photobleaching-related phenomenon is photochromism, which we systematically characterised for all FP-T2A-FP constructs. The control FPs in the T2A constructions unlikely affect photochromism as this behaviour is determined by measuring only the fluorescence of the FP of interest. Besides, we did not observe any photoswitching of mTq2 or mCherry, which could affect the fluorescence readout of the FP of interest. We show that photochromism only occurs when exciting with a higher-energy wavelength than the optimum wavelength of the specific FP. It could be that this light energy changes the photochrome conformation and thereby changes the fluorescent state the FP is in, as suggested before (Pletnev et al., 2008). Additionally, the specific wavelength used does not largely affect photochromic behaviour (Table S1). Interestingly, various photochromic FPs already start in a dark-state and are immediately pushed into a fluorescent state by excitation lights of shorter wavelengths than the optimal wavelength for the FP. This causes photoactivation, which can make cells more than 3 times brighter than their starting brightness. This has large implications as experiments with photochromic FPs can generate experimental bias due to photochromism.

However, photochromism can also be used as an advantage. Using photochromic FPs under dual-excitation for long timelapse experiments reduces bleaching, making these FPs highly photostable (Fig. 3C). Still, for other experiments, photochromic FPs should be used with care in any organism as they can generate unnoticed quantitative artefacts which can result in wrongly drawn conclusions. With the supplied model, we could fit all bleaching kinetics obtained in the present study (Fig S3). Therefore, we expect that our set of equations that make up the model can be used to quantify, predict and correct for complex bleaching kinetics using other setups and conditions as well. We recommend to use the model when experiments with photochromic FPs are necessary and unevitable to ensure correct quantification of fluorescent signals.

Lastly, to our knowledge, the pH sensitivity was determined for the first time *in vivo* for all FPs. We found that the pKa values of FPs can differ up to more than 0.5 pH point between *in vivo* and *in vitro* for several FPs (Table S1). Therefore, the performance of an FP in a pH dynamic or low-pH environment should be assessed *in vivo* and not *in vitro* before performing experiments to rule out any pH bias. In *S. cerevisiae,* pH is tightly controlled but very dynamic and even implied as second messenger (Dodd and Kralj, 2017; Orij et al., 2012, 2009), and so experiments that affect the physiology of yeast should preferably be studied with pH-insensitive FPs. Which factors specifically affect pH sensitivity *in vivo* is not known, although it is known that ions such as Cl^−^ affect the pH curves (Griesbeck et al., 2001). We believe that also other cellular components such as adenosines, NAD and NADP can affect these curves as they also affect CFP-YFP FRET ratios (Moussa et al., 2014). Interestingly, although the CFPs were still pH-robust with pKas lower than 5 and a Hill coefficient of approximately 0.5, we found them more pH-sensitive than determined *in vitro.* For GFPs, mNeongreen is the best performer although it is still rather pH sensitive. Yet, a pH-robust GFP (pKa of 3.4) was developed recently (Shinoda et al., 2018).

Based on the measured FP traits, we created yeast codon-optimized FP of the best performing FPs and found -to our surprise-that codon-optimization affected brightness even though the primary sequence is unaffected. The practical FP brightness is the product of the quantum yield, extinction coefficient and the amount of matured FPs. Codon optimization probably does not affect the quantum yield and absorption as these are determined by the protein structure itself. Yet, the folding, functioning and degradation of proteins can be affected by codon optimization and this can affect the practical brightness of FPs (Yu et al., 2015; Zhou et al., 2013). This is the case for all yFPs: ymVenus, ymTq2, ymNeongreen show increased brightness whereas ymScarletI, ytdTomato and ymYPET show decreased brightness. Therefore, codon-optimization should be performed with precaution and the codon-optimized proteins should be checked for proper functioning before usage. Although ytdTomato and ymScarletI have decreased brightness, they have improved FP expression, making the absolute signal comparable to their non-optimized variants (Figs 5A & 5C). However, the decreased practical brightness can be caused by increased degradation of the FPs. Therefore, both the codon-optimized as the original variant of tdTomato and mScarletI should be used carefully for endogenous tagging as they can affect protein levels, either by inducing low expression (for the original FP variants) or by increased protein degradation (for the codon-optimized variants). The non-optimized variants of ytdTomato and ymScarletI can be used for plasmid-based overexpression as these FPs are brighter. One important note is the size of tdTomato, which is twice the size of the other FPs. This can be of importance when using it for protein tagging. The large decrease in ymYPET brightness compared to YPET is probably also due to the altered protein structure to make it monomeric. Yet, ymVenus is not monomeric, making ymYPET still the preferred YFP with an relatively high brightness, high pH-insensitivity and high photostability for the YFP class.

Finally, we show that the developed tagging vectors can be conveniently used to tag proteins endogenously in yeast. We used the tagging vectors to tag FBPase with yoeGFP, yosfGFP or the novel ymNeongreen. The use of ymNeongreen increases the dynamic range of a fluorescent-based flow cytometry experiment (Fig 8). By using ymNeongreen instead of yoeGFP or yosfGFP, we were able to fully separate populations with an induced or uninduced FBPase. This shows the importance of performing experiments with the correct –and best performing–FPs, even compared to “gold standard FPs” such as eGFP.

In conclusion, we show that FP characteristics are context dependent and should be characterised per specific organism *in vivo.* The yFP palette that we generated consists of the best performing FPs for every spectral class in yeast to date in terms of brightness, photostability, photochromism and pH robustness. The use of the yFPs opens new experimental possibilities and enhances fluorescent-based readouts compared to the previously codon-optimized FPs (Lee et al., 2013; Sheff and Thorn, 2004). We anticipate that our results will also be more representative for FP perfomance in organisms grown at low temperatures (e.g. zebrafish, Caenorhabditis elegans, Drosophila melanogaster and Arabidopsis) than mammalian-based characterisations. Our provided FP properties can be taken into consideration to improve experimental succes and reliability.

## Acknowledgements

We thank Paula Martínez Román (Erasmus scholarship BSc student at Vrije Universiteit Amsterdam) and Jip Wulffelé (BSc student at Vrije Universiteit Amsterdam) for constructing various FP constructs and assisting with experiments. We thank Rick Nijhout for constructing pFA6a-link-ymNeongreen SpHis5. We thank Laura van Weeren and Marieke Mastop (Molecular Cytology, University of Amsterdam) for constructing the plasmids for co-expressing proteins with the T2A sequence in mammalian cells.

This work was supported in part by the NWO VICI grant 865.14.005 and the F10.001.03 BE Basic project under the TKIEB01003 AMBIC program.

